# Integrated metabolomics and proteomics of symptomatic and early pre-symptomatic states of colitis

**DOI:** 10.1101/2020.03.22.002196

**Authors:** Elee Shimshoni, Veronica Ghini, Inna Solomonov, Claudio Luchinat, Irit Sagi, Paola Turano

**Affiliations:** Department of Biological Regulation, Weizmann Institute of Science, Rehovot, Israel; Center of Magnetic Resonance (CERM), University of Florence, Florence, Italy; Department of Chemistry, University of Florence, Florence, Italy

**Author notes:** Authors contributed equally (Elee Shimshoni, Veronica Ghini).

**Keywords:** Inflammatory bowel disease, metabolomics, proteomics, colon tissues, fecal extracts

## Abstract

Two murine models for colitis were used to study multi-level changes and derive molecular signatures of colitis onset and development. By combining metabolomics data on tissues and fecal extracts with proteomics data on tissues, we provide a comprehensive picture of the metabolic profile of acute and chronic states of the disease, and most importantly, of two early pre-symptomatic states. We show that, increased anaerobic glycolysis, accompanied by altered TCA cycle and oxidative phosphorylation, associates with inflammation-induced hypoxia taking place in colon tissues. We also demonstrate significant changes in the metabolomic profiles of fecal extracts in different colitis states, most likely associated with the dysbiosis characteristic of colitis, as well as the dysregulated tissue metabolism. Most remarkably, strong and distinctive tissue and fecal metabolomic signatures can be detected before onset of symptoms. These results highlight the diagnostic potential of global metabolomics for inflammatory diseases.

## Introduction

Inflammatory bowel disease (IBD), which primarily includes ulcerative colitis (UC) and Crohn’s disease (CD), is a progressive, chronic and relapsing condition. This debilitating disease is steadily becoming a worldwide medical concern, with increasing prevalence and incidence in both industrialized and developing countries (Molodecky *et al*, 2012). For this reason, detection of molecular signatures, at the tissue level, that aid early diagnosis and prediction of flares is critical for disease management.

IBD is a multifactorial disorder, and genes associated with predisposition to IBD encode critical players in the innate immune response, inflammation, autophagy, and epithelial barrier integrity (Shah, 2016). Moreover, epithelial oxygen tension is a common feature of intestinal inflammation in IBD (Shah, 2016). While physiological hypoxia is localized to epithelial cells adjacent to the anoxic lumen, in colitis hypoxic staining is observed throughout the mucosa, likely because of the inflammation-driven enhanced oxygen consumption of intestinal epithelial cells and the decreased oxygen availability to inflamed areas due to vasculitis. Under healthy conditions, epithelial hypoxia limits the amount of oxygen emanating from the mucosal surface, which helps maintain anaerobiosis in the intestinal lumen and ensures the colonic microbiota is dominated by obligate anaerobic bacteria, that convert fiber into fermentation products (Litvak *et al*, 2018). Microbiota-derived short-chain fatty-acids (SCFA; particularly acetate, propionate and butyrate) regulate oxygen consumption in intestinal epithelial cells (Kelly *et al*, 2015), influence the host immune response and can promote interleukin-10 (IL-10) production in T-helper type 1 (Th1) cells by taking part in a dynamic host-microbiome network (Sun *et al*, 2018). Through this mechanism, the colonic epithelium shapes the microbiota to be beneficial, thereby maintaining gut homeostasis (Albenberg *et al*, 2014). Dysbiosis in the colon, such as in individuals suffering from IBD, is commonly associated with an expansion of facultative anaerobic bacteria (Lloyd-Price *et al*, 2019).

IL-10 is a key anti-inflammatory cytokine produced by activated immune cells, which inhibits the activity of inflammatory (Ip *et al*, 2017). Morover, the gastrointestinal tract contains the largest pool of macrophages in the body, which are key players in maintaining intestinal homeostasis (Bain & Schridde, 2018). It has been reported (Ip *et al*, 2017) that under conditions of IL-10 deficiency, macrophagesare characterized by an exaggerated glycolysis and loss of mitochondrial fitness. Similarly, other authors have reported that pro-inflammatory macrophages are more glycolytic, produce more ROS, accumulate succinate and have a suppressed oxidative phosphorylation with respect to resting macrophages (O’Neill & Pearce, 2016). In turn, the levels of succinate affect the activity of hypoxia-inducible factor 1 (HIF-1), a key transcription factor in the expression of pro-inflammatory genes (Tannahill *et al*, 2013). In activated macrophages (M1 subtype) an increased flux through the aspartate–arginosuccinate shunt feeds the Krebs cycle at fumarate, replenishing the cycle after the break point that occurs in M1 macrophages at the enzyme succinate dehydrogenase (SDH) (Mills *et al*, 2017).

Here, we utilize two common murine models for IBD – the acute dextran sodium sulfate (DSS)-induced colitis model (Okayasu *et al*, 1990), and the chronic piroxicam-accelerated colitis (PAC) model in interleukin(IL)-10^−/−^ mice (Berg *et al*, 2002a) for studying multi-level changes in metabolic profiles over the course of colitis development. We examine both types of disease states (acute and chronic), and most importantly, we examine two early states, from each model, in which there are no detectable symptoms, and therefore can be regarded as “pre-symptomatic”. Nuclear magnetic resonance (NMR) metabolomics data on tissues and fecal extracts and liquid chromatography–tandem mass spectrometry (LC-MS/MS)-based proteomics data on tissues were combined to derive the molecular signature of disease onset and progression. From these data, we demonstrate i) how metabolomic profiling based on high-resolution magic angle spinning (HR-MAS) ^1^H NMR of tissues can monitor the global metabolic shift associated with hypoxia also at early, preclinical states; ii) how tissue proteomics nicely complements the NMR findings; iii) how solution ^1^H NMR profiling of fecal extracts reveals pre-symptomatic states showcasing it potential in future applications as a non-invasive early diagnosis tool.

## Results and Discussion

### Clinical characterization and timeline of murine colitis models

By using two different colitis murine models we examine acute and chronic states, and most importantly, we examine also early “pre-symptomatic” states. In this subsection, we endoscopically and histologically define these states and disease models, in a similar manner to our work in (Shimshoni *et al*, 2019).

Acute colitis was induced in wild type (WT) C57BL/6 mice via addition of 1.25% DSS into the drinking water for seven days (**Fig 1A**). Disease progression was monitored by endoscopic evaluation and histological analysis of colonic tissue at different time points (**Fig S1**). Inflammation degree was scored during endoscopic examination according to five parameters: mucosal transparency, mucosal wall granularity, stool consistency, blood vessel deformation and fibrin appearance in the colon lumen. Each parameter was given a score between 0-3 (Becker *et al*, 2006), for an overall score of 0-15. An overall score of 0-4 was defined as “healthy”; 5-7 was defined as “mildly inflamed”; 8-11 as “inflamed”; and 12-15 as “severely inflamed”. Colon inflammation under these conditions peaks between days 7-10 (**Fig 1A**), as can be detected by endoscopy (**Fig S1A**). At day 10 all mice present with the two highest categories of colon inflammation. Importantly, clinical symptoms begin to appear after day 5. On day 4 animals are defined by endoscopic evaluation as “healthy” (**Fig S1A**) and seldom show mild immune cell infiltration as assessed by histological analysis (**Fig S1B**). Therefore, we regard day 4 as the “pre-symptomatic” state.

**Figure 1.**
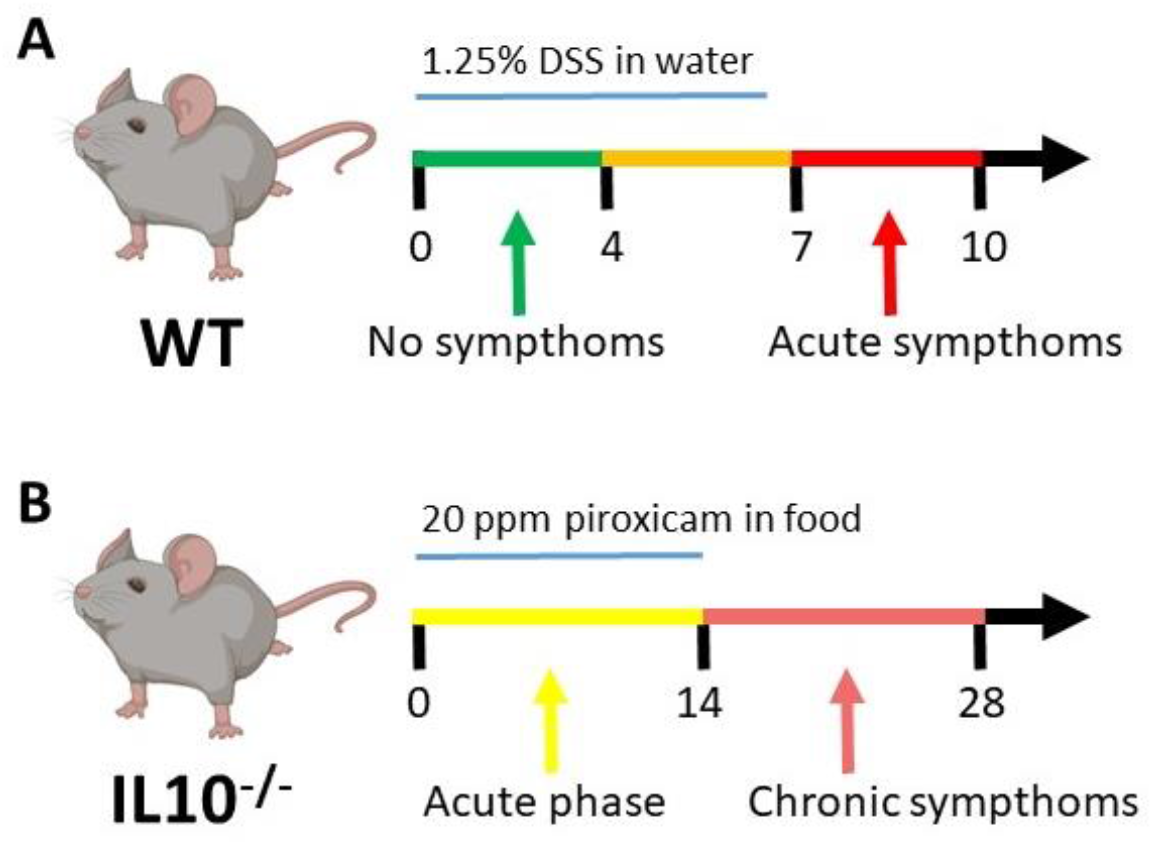
Course of disease for the two murine colitis mode. A. Timelines, acute model: Wild-type (WT) C57BL/6 mice, exposed to 1.25% (w/v) DSS in the drinking water for seven days; clinical and endoscopic symptoms peak on day 10. B. Timelines, chronic model: C57BL/6 IL-10^−/−^ mice, exposed to 200 ppm piroxicam in the food for 14 days; clinical and endoscopic symptoms develop over the course of these 14 days and persist as chronic inflammation for at least 14 days.

To gain insight on chronic intestinal inflammation, we used C57BL/6 IL-10−/− mice that spontaneously develop chronic colon inflammation (Kühn *et al*, 1993). This inflammation is not chemically induced, but rather is considered to be a result of a dysregulated immune system, which more resembles the clinical manifestation of chronic UC and CD patients and mimics a monogenic form of IBD (Glocker *et al*, 2009). In order to synchronize and accelerate the onset of chronic colitis, mice were exposed to 200 ppm piroxicam in their food for 14 days (**Fig 1B**) (Berg *et al*, 2002a). The result of this exposure was the development of IBD-like symptoms, as can be detected and scored using colonoscopy in the same manner as for the well-established DSS-induced model (**Fig S1A**). Histopathology in this model shows prominent immune cell infiltration into the mucosa, but with little ulceration or colonic crypt abolition (**Fig S1B**). The two models allow us not only to study different kinds of colitis, the transient acute colitis and the persistent chronic colitis; but also lead us to “zoom in” onto two pre-symptomatic states, or inflammation-prone states – day 4 of the DSS model and the healthy naïve IL-10^−/−^ mouse, by monitoring disease progression on a daily basis. With the described experimental procedures, we obtained five groups of mice for further analysis: 1) Healthy WT; 2) Day 4 of DSS (pre-symptomatic); 3) Day 10 of DSS (acute colitis); 4) Healthy IL-10^−/−^ (pre-symptomatic); and 5) Ill IL-10^−/−^ (chronic colitis).

### Colitis leads to a shift towards anaerobic metabolism in the colon

We set out to analyze metabolic changes that take place over the course of colitis, at the chosen five states mentioned above. To this end, the colon tissues were analyzed by untargeted HR-MAS ^1^H NMR metabolomics, untargeted LC/MS-MS proteomics and real-time PCR for a subset of genes involved in glycolysis.

^1^H NMR spectra were acquired on mouse colon tissues with the final aim of identifying the metabolomic fingerprint of control, pre-symptomatic and symptomatic states. The spectra demonstrated reproducibility within each of the five groups. The metabolomic fingerprint in all spectra is made up of both broad and sharp features accounting for more than 130 resonances with different relative intensities within the 5 groups (**Fig S2**). This untargeted NMR approach (Takis *et al*, 2018; Vignoli *et al*, 2019) revealed that each state in both models has a distinct metabolic profile (**Fig S3**). The characteristic metabolomic profile of each state can be distinguished when all five states are considered together, with a 5-group 72% discrimination accuracy, according to a PCA-CA model (**Fig 2A**).

**Figure 2.**
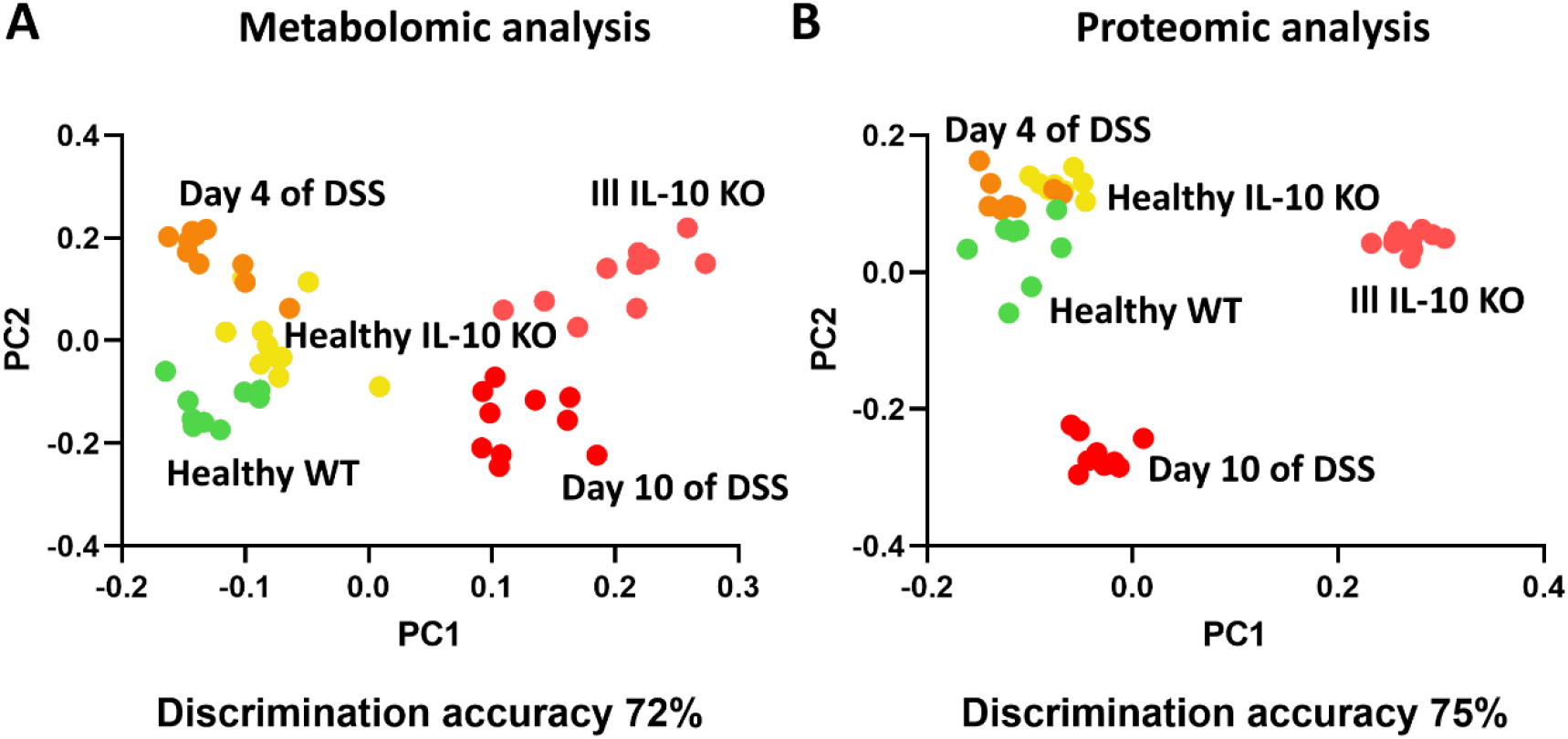
Metabolomic and proteomic profiles of different colitis states. A. PCA-CA analysis of NMR data. The PCA-CA score plots reveal different profiles of the five states in colitis, with a 5-group discrimination accuracy of 72%. B. PCA-CA analysis of LC-MS/MS proteomic data. The PCA-CA score plots reveal different profiles of the five states in colitis, with a 5-group discrimination accuracy of 75% for proteomics. In the plots each dots represents a different sample and each color a different group of mice: green dots, healthy WT (WT-HC) mice; orange dots, WT-D4; yellow dots, IL10-HC; red dots, WT-D10; pink dots, IL10-ILL.

LC/MS-MS proteomics was based on a dataset of 2045 non-redundant proteins expressed in all the five states, deriving from a previous work by some of us (Shimshoni *et al*, 2019). Out of them, 425 proteins involved in metabolic processes were identified using the Human Metabolic Atlas database (http://www.metabolicatlas.org). Considering this subset of proteins, the five different tissue states display differentiating proteomic profiles, with 5-group 75% discrimination accuracy, according to a PCA-CA model. (**Fig 2B**). Both analytical platforms well discriminate the symptomatic states from the corresponding controls. Importantly, pre-symptomatic states (day 4 of the DSS model and healthy IL-10^−/−^) can be discriminated from healthy WT samples despite no macroscopic inflammation is observed.

The NMR spectra analyses lead to the identification of 22 metabolites (**Table S1**); several of them have significantly different levels in the various groups. Among those mainly responsible for the discrimination between healthy and pathological tissues, there are products of the glucose metabolism; in both chronic and acutely inflamed tissues, a significant increase in glucose levels was accompanied by a significant increase in lactate levels (**Fig 3A)**. The observed anaerobic shift in colitis is consistent with the hypoxic conditions that develop during inflammation (Shah, 2016; Giatromanolaki *et al*, 2003; Taylor & Colgan, 2007). Consistently, a western blot analysis for hypoxia-induced factor (HIF)-1α on tissue extracts indicated high levels of HIF1α only in inflamed tissues (**Fig 3A**). Regarding glycolytic proteins, proteomics and mRNA data do not show clear trends, and the interpretation is complicated by the fact that glycolysis is mainly regulated by post-translational modification and feedback mechanisms rather than by changes in the enzyme levels (Berg *et al*, 2002b), (**Fig S4, Table S2**).

**Figure 3.**
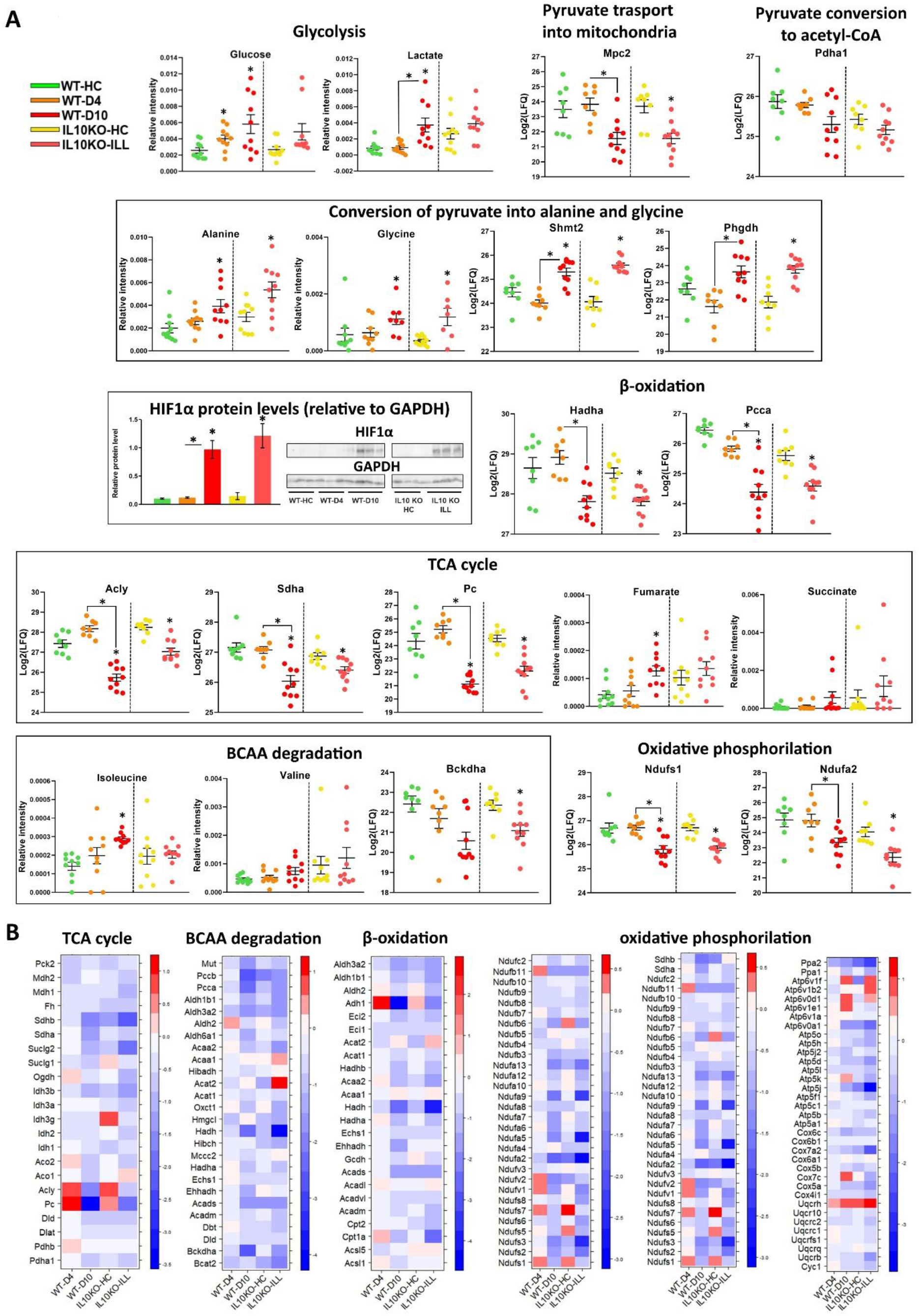
Metabolomic and proteomic analysis of colon tissues. A. PCA-CA analysis of NMR data. The PCA-CA score plots reveal different profiles of the five states in colitis, with a 5-group discrimination accuracy of 72%. B. PCA-CA analysis of LC-MS/MS proteomic data. The PCA-CA score plots reveal different profiles of the five states in colitis, with a 5-group discrimination accuracy of 75% for proteomics. C. Dot plots of relative concentration levels of selected proteins and metabolites in the five colitis states and Western blots for hypoxia-induced factor (HIF)1α. Proteins and metabolites are grouped according to the metabolic process in which they are involved. Asterisks indicate significance relative to corresponding healthy state (either WT or IL10KO), unless otherwise indicated by bars. *p<0.05. D. Heat maps for relative mean abundance (in log2(LFQ) compared to healthy WT) of proteins involved in different metabolic processes. Downregulated / upregulated proteins are shown using blue / red color coding. In the plots each dots represents a different sample and each color a different group of mice: green dots, healthy WT (WT-HC) mice; orange dots, WT-D4; yellow dots, IL10-HC; red dots, WT-D10; pink dots, IL10-ILL.

The analysis of other metabolites, directly or indirectly, produced by glycolysis provided further information. Pyruvate levels remain below detection in our metabolomic data but some hints on the fate of this metabolite can be derived indirectly. Besides being reduced to lactate, pyruvate can be transaminated to produce alanine, whose levels are also increased both in samples from mice on day 10 of the DSS model and from ill IL-10^−/−^ mice, compared to their healthy counterparts (**Fig 3A**). A significant portion of pyruvate can also be diverted toward glycine biosynthesis by phoshphoglycerate dehydrogenase (Phgdh) and serine hydroxymethyltransferase (Shmt2). Consistently, glycine, Phgdh, and Shmt2 levels are increased in day 10 and ill IL 10^−/−^ samples (**Fig 3A**).

Our metabolomics and proteomics data also highlight alterations of the TCA cycle (**Fig 3A and B, Table S2**). Pyruvate can enter in the mitochondria to fuel the TCA cycle, being transformed into acetyl-CoA. Acetyl-CoA remains below the NMR detection limit, but we observed a reduction in the levels of the proteins involved, not only in pyruvate transport into the mitochondria (i.e., mitochondrial pyruvate carrier 2, Mpc2) (**Fig 3A**), but also in its conversion into acetyl-CoA (downregulation of the proteins of the Pyruvate Dehydrogenase Complex, (PDC), i.e., pyruvate dehydrogenase subunit 1, Pdha1, dihydrolipoyl transacetylase, Dlat; dihydrolipoamide dehydrogenase, Dld) (**Fig 3A**).

Along the same lines, most of the proteins involved in the TCA cycle itself (e.g., ATP-citrate lyase, Acly; succinate dehydrogenase complex subunit A, Sdha; pyruvate carboxylase, Pc) are less abundant in both pathological mouse tissues (**Fig 3A and B**).

Analogously, the metabolomic profiles highlighted an increase in the levels of the TCA cycle intermediates succinate and fumarate (**Fig 3A**), which accumulate due to inefficient TCA metabolism (Bernacchioni *et al*, 2017; Caracausi *et al*, 2018).

In addition to glycolysis-derived pyruvate, fatty acids and amino acids can supply substrates to the TCA cycle. The breakdown of fatty acids (β-oxidation) in the mitochondria generates acetyl-CoA. Our proteomic results show that most of the enzymes taking part in the β-oxidation pathway display lower levels in chronic and acute colitis (e.g., hydroxyacyl-CoA dehydrogenase trifunctional multienzyme complex subunit α, Hadha; propionyl-CoA carboxylase subunit α, Pcca) (**Fig 3A and B, Table S2**), thus further reducing the cellular content of acetyl-CoA available to sustain the TCA cycle.

Also branched chain amino acids (BCAA), isoleucine, valine and leucine, can be converted into acetyl-CoA and other organic molecules that enter the TCA cycle. Accordingly, we detected a strong reduction of proteins involved in the BCAA degradation pathway by branched chain keto acid dehydrogenase E1, α polypeptide, (Bckdha) (**Fig 3A and B, Table S2**). These data correlate well with an increasing trend in the levels of isoleucine and valine in pathological tissues (**Fig 3A**).

Finally, and in agreement with all previous results, acute and chronic colitis tissues are characterized by a reduction in the protein involved in the oxidative phosphorylation such as some of the components of the Complex I – e.g., NADH:ubiquinone oxidoreductase core subunits s1, s2, s3, s5, s6, s7, v1, a2 and a6 (Ndufs1, Ndufs2, Ndufs3, Ndufs5, Ndufs6, Ndufs7,Ndufv1, Ndufa2, Ndufa6) (**Fig 3A** and **B, Table S2**).

Hence, by combining the two “omic” analyses, we were able to achieve a comprehensive picture of the metabolic profile of the intestinal tissues at each state, both acute and chronic. Increased anaerobic glycolysis is accompanied by an altered TCA cycle due to depressed acetyl-CoA production and consequent accumulation of fumarate and succinate; altered oxidative phosphorylation is suggested by reduced levels of Complex-I proteins (**Fig 4**). These results are in line with the metabolic characteristics of activated M1 intestinal macrophages (Mills *et al*, 2017).

**Figure 4.**
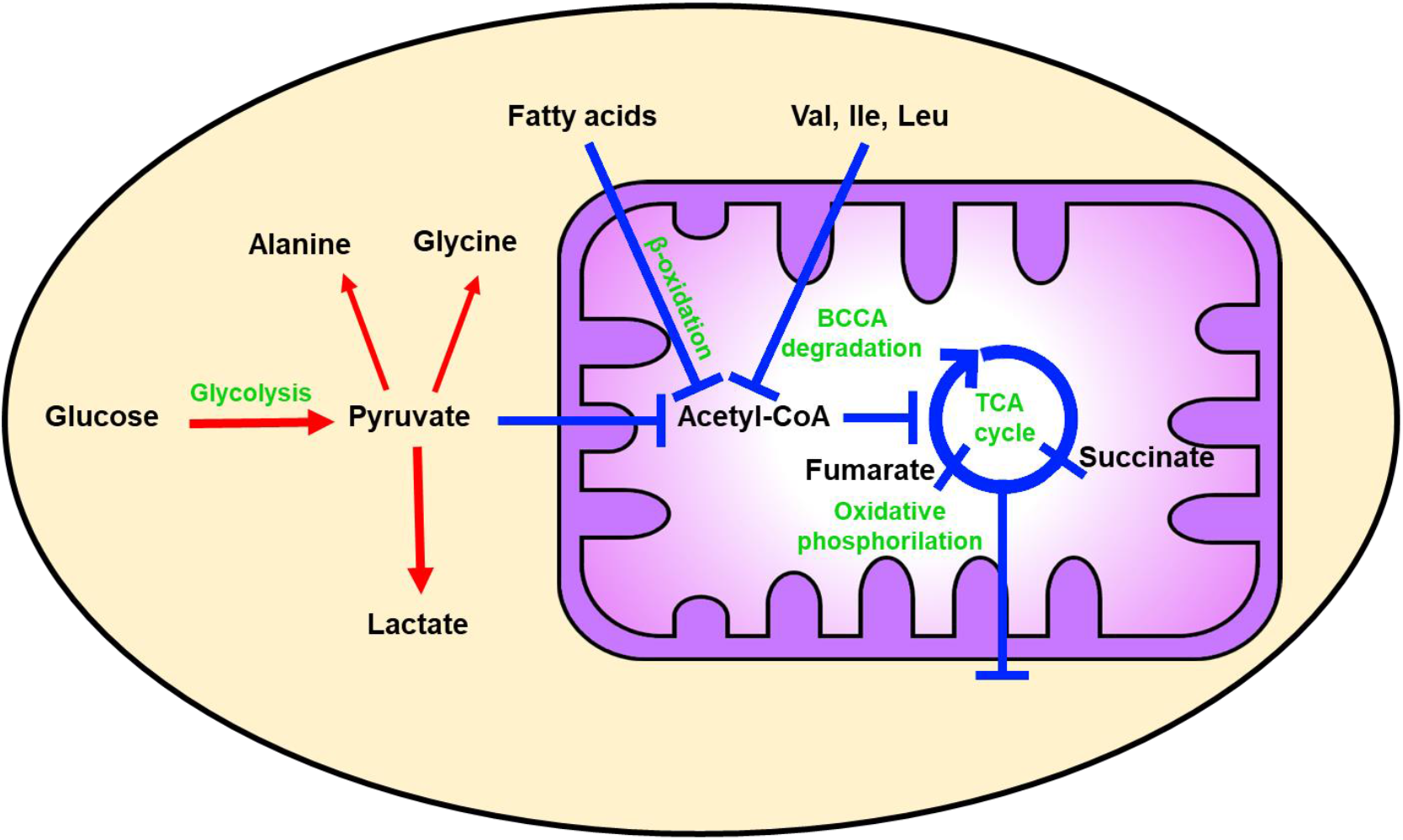
Colitis leads to a metabolic shift in tissues. The major metabolic processes that are differentially regulated in both acute and chronic colitis are indicated. Upregulated processes are described by red arrows and downregulated processes by blue bars.

### Pre-symptomatic metabolic profiles are shared across models

Another key finding of this study is that pre-symptomatic states have a characteristic metabolic signature. Close investigation of our analysis of the DSS colitis model reveals that the pre-symptomatic state, day 4, does not simply represent a progressive shift towards colitis, with some proteins (e.g., Pfkl, Acly and Pc) and transcript (e.g., Pflk and Pgk1) levels displaying an irregular trend. To assess the metabolic and proteomic similarity between the two pre-symptomatic states, we generated a metabolomic and a proteomic PCA-CA model on all samples except those of day 4 of the DSS model. When predicting day 4 of DSS samples on both the two models they were mostly predicted as belonging to the class of healthy IL-10^−/−^ samples (with 80% accuracy for the metabolomic model and 88% accuracy for the proteomic model) (**Fig 5**). These findings demonstrate the convergence of the two “paths” towards inflammation, whether chronic or acute, with similar pre-symptomatic states.

**Figure 5.**
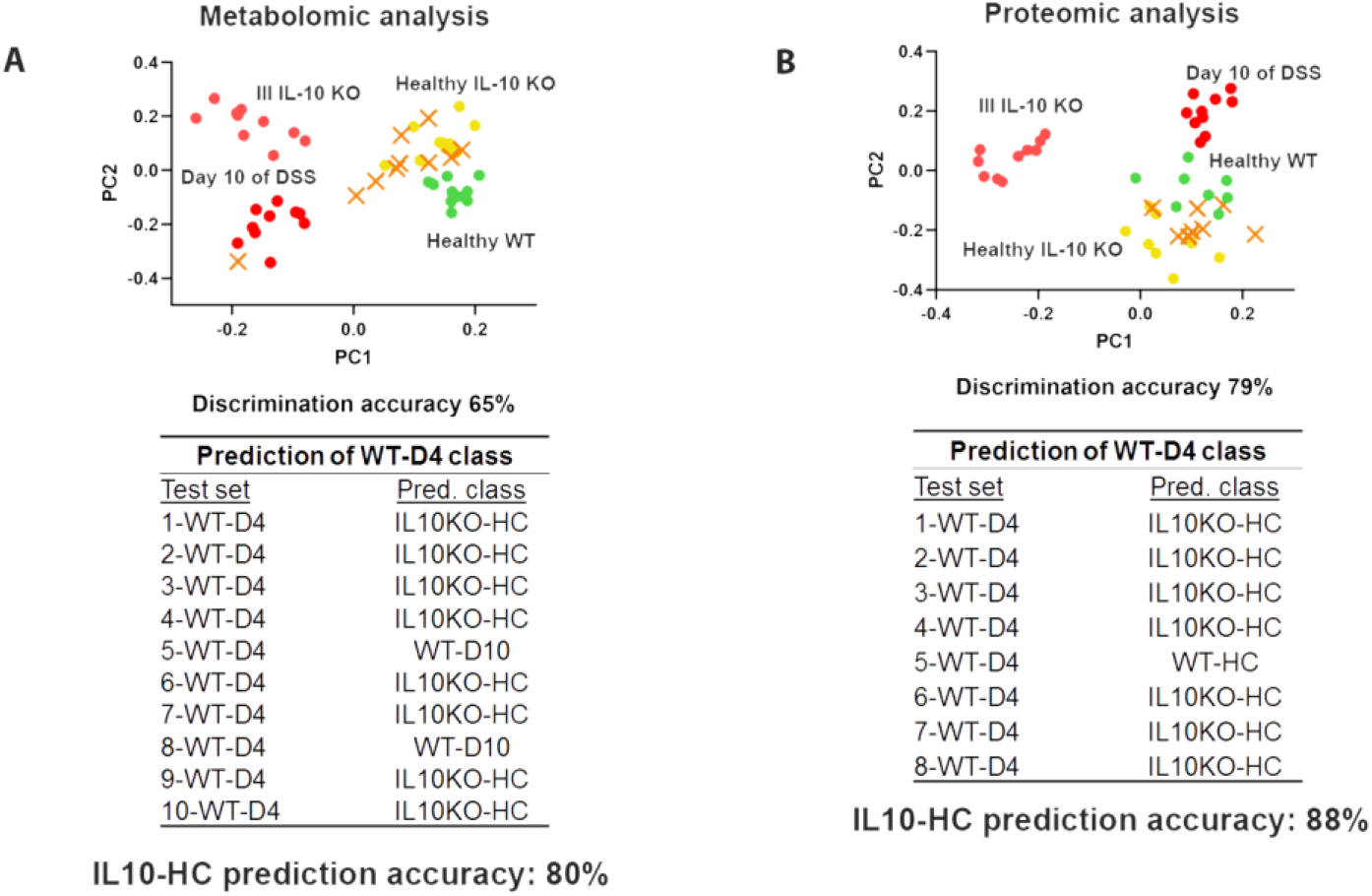
Pre-symptomatic metabolic profiles of tissues are shared across models. A. PCA-CA prediction of WT-D4 group (orange crosses) using a training set composed of the other four groups according to metabolomics. WT-D4 samples are mainly predicted as the other pre-symptomatic state – IL10^−/−^HC, with a prediction accuracy of 80%. In particular, only two samples (5-WT-D4 and 8-WT-D4) are predicted as more advanced (WT-D10). B. PCA-CA prediction of WT-D4 group (orange crosses) using a training set composed of the other four groups according to proteomics. WT-D4 samples are mainly predicted as the other pre-symptomatic state – IL10^−/−^HC, with a prediction accuracy of 88%. In particular, only one sample (5-WT-D4) is predicted as les advanced (WT-HC). In the score plots, each symbol represents a different sample and each color a different group of mice: green dots, WT-HC; orange crosses, WT-D4; yellow dots, IL10-HC; red dots WT-D10; pink dots, IL10-ILL.

### Fecal metabolomic analysis can serve as a diagnostic and prognostic tool for intestinal inflammation

Dysbiosis has been implicated as a hallmark and important player in IBD (Tamboli *et al*, 2004; Lloyd-Price *et al*, 2019), we sought to examine whether fecal metabolomics can be a good indicator of inflammation, and especially of pre-clinical states. To this end, we performed metabolomic analysis on fecal extracts corresponding to the five tissue states discussed above (healthy WT, day 4 of DSS, day 10 of DSS, healthy IL-10^−/−^ and chronically ill IL-10^−/−^). An untargeted metabolomic fingerprinting approach demonstrated that each state has its own unique fecal metabolomic profile, as shown by the results of the PCA-CA analysis (**Fig 6A**). Among the five groups, our analysis provided a discrimination accuracy value of 93% (**Fig 6A**). Testing day 4 samples of the DSS model according to a PCA-CA model constructed for the other four groups reveals, also in this analysis, the similarity among the pre-symptomatic states – with an 89% prediction accuracy for day 4 of DSS as healthy IL-10^−/−^ samples (**Fig 6B**). In the NMR spectra of fecal extracts 35 metabolites (**Table S3**) were identified and quantified; they belong to different chemical groups, such as fatty acids, amino acids and alcohols. In **Fig 6C**, the results of univariate analysis on selected metabolites are presented.

**Figure 6.**
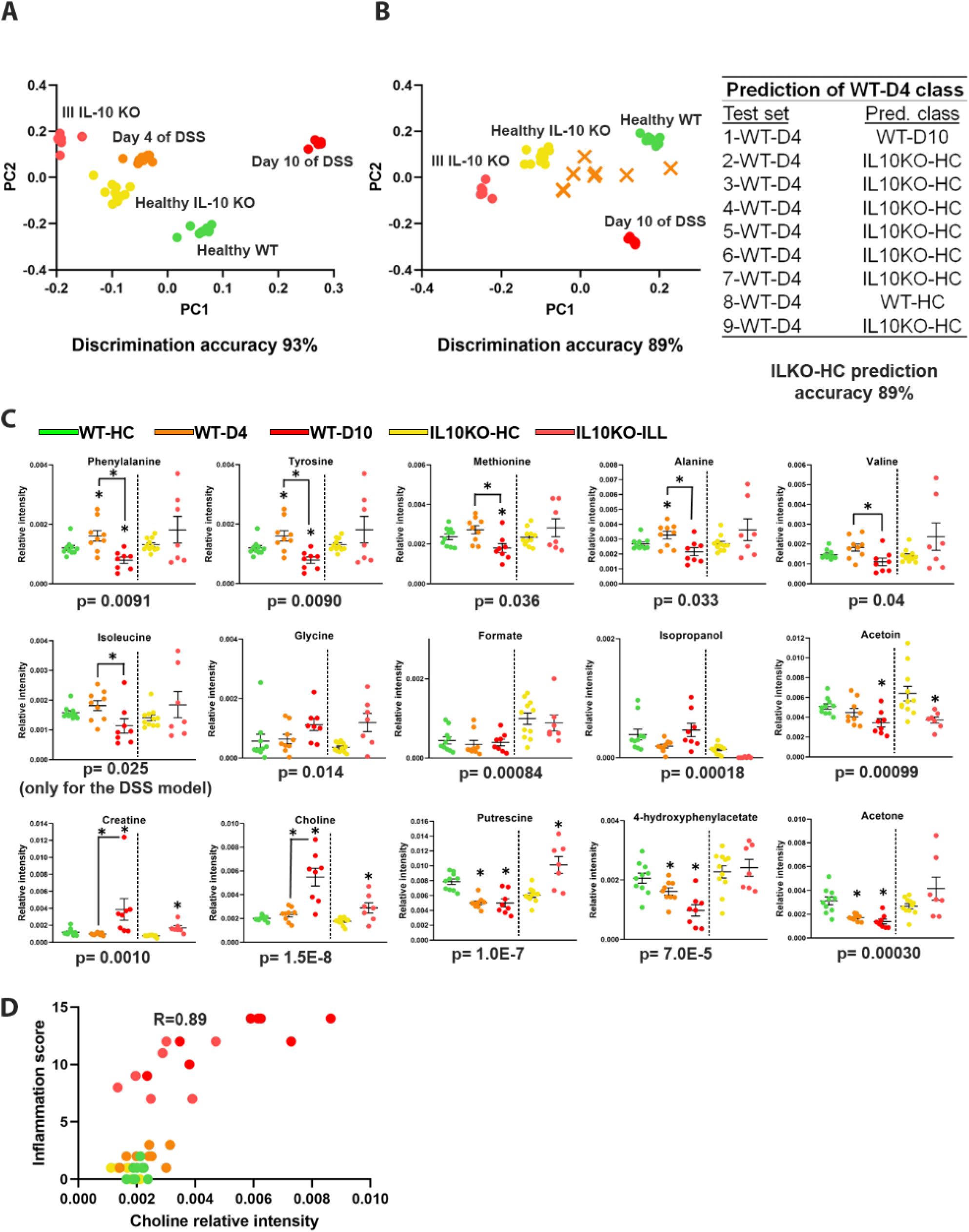
Fecal metabolomic analysis. A. PCA-CA analysis of NMR-based metabolomic data of faecal extracts. The PCA-CA score plot reveals that feces have significantly different metabolomic profiles in the five colitis states of the two models, with a 5-group discrimination accuracy of 93%. B. PCA-CA prediction of WT-D4 group (orange crosses) using a training set composed of the other four groups (4-group discrimination accuracy 89%). WT-D4 samples are mainly predicted as the other pre-symptomatic state – IL10-HC, with a prediction accuracy of 89%. In particular, only one sample (1-WT-D4) is predicted as more advanced (WT-D10) and another one as less advanced (WT-HC). C. Dot plots of relative concentration levels of selected metabolites in the five states. Asterisks indicate significance relative to corresponding healthy state (either WT or IL10KO), unless otherwise indicated by bars. *p<0.05. D. Plot depicting the polychoric correlation between fecal choline levels and endoscopic score (R=0.89). In all plots, each symbol represents a different sample and each color a different group of mice: green dots, WT-HC; orange dots, WT-D4; yellow dots, IL10-HC; red dots, WT-D10; pink dots, IL10-ILL.

Among the quantified metabolites there are butyrate, propionate and acetate, which are the main SCFA produced during fermentation by gut bacteria (McNeil *et al*, 1978; Topping & Clifton, 2001). Interestingly, the levels of the metabolites involved in glucose metabolism, such as glucose, lactate, pyruvate and fumarate, are not significantly altered in the samples from chronic and acutely inflamed mice (**Fig S5**), indicating that the metabolome of feces does not directly reflect that of tissues, but probably the crosstalk between the microbiome and the host.

SCFA are the major nutrients produced by bacterial fermentation of carbohydrates. Here the fecal butyrate, propionate and acetate were slightly increased, albeit not significantly, in chronically ill IL-10^−/−^ with respect to healthy IL-10^−/−^, while no significant trend was identified comparing healthy WT and day 10 of DSS (**Fig S6**). Interestingly, an increase of these metabolites was detected when comparing day 4 of DSS with respect to healthy WT. Butyrate, propionate and acetate higher levels in day 4 of DSS where similar to the levels of these metabolites in healthy IL-10^−/−^, again supporting the similarity between the pre-symptomatic states. However, these findings seem to point to a different behavior in murine models compared to the human disease, as, SCFA levels have been reported to not change significantly in fecal samples of DSS-treated mice (Osaka *et al*, 2017), whereas reduced levels of SCFA in feces of IBD patients have been reported and linked to a shift in the composition and in the metabolic activity of intestinal microbiota, specifically in the reduction in butyrate-producing bacterial groups (De Preter *et al*, 2015; Marchesi *et al*, 2007; Bjerrum *et al*, 2015).

SCFA are mostly absorbed in the colon while less than 5% are excreted in the feces. Our results may reflect a reduced absorption/uptake and oxidation of SCFA during inflammation, indicated by the downregulation of some enzymes involved in butyrate metabolism/oxidation such as EHHADH, HADHA, PDHA1, ACAT1, ACAT2, ALDH2, and ACADS in acute and chronic inflamed tissues (**Fig 3**). Genes encoding butyrate uptake and oxidation were also found to be down-regulated in inflamed mucosa of UC patients (Parada Venegas *et al*, 2019; De Preter *et al*, 2012). Hence, these results show that inflammation is tightly linked to the inhibition of genes related to SCFAs uptake and metabolism.

We also observed a peculiar trend for the amino acids valine, isoleucine, alanine, methionine, tyrosine and phenylalanine. In the acute colitis model, their levels increase on going from healthy WT to day 4 of DSS, and then decrease on day 10 of DSS; while in the chronic colitis model their levels do not change (**Fig 6C**). No changes in either model are found in the amino acid glutamate (**Fig 65**). On the contrary, glycine shows a regular trend, significantly increasing in its levels in fecal samples from both inflamed states, with respect to their healthy controls. It is known that amino acids play significant roles in intestinal inflammation (He *et al*, 2018), being involved in multiple signaling mechanisms related to intestinal inflammation. For example, isoleucine and valine activate the GCN2 pathway; methionine and glycine inhibit NF-κB signaling pathway; and tyrosine, valine and phenylalanine activate CaSR. No specific role is yet reported for alanine. Whether these differences reflect decreased absorption or increased loss by inflamed intestines needs to be elucidated, but is consistent with previous findings on pediatric IBD patients (Bosch *et al*, 2018).

Creatine and choline display the same behavior described above for glycine (**Fig 6C**). The increase in creatine excretion in both pathological states may be attributed to muscle dysfunction leading to reduced ATP consumption via creatine phosphorylation. 4-hydroxyphenylacetate decreases only in samples from mice with acute inflammation, while acetoin decreases in both colitis states. The levels of acetone and putrescine decrease in samples from mice with acute inflammation while they increase in those with the chronic disease (**Fig 6C**). Most notably, choline, which is a component of many biological molecules, like acetylcholine and lipids, has a strong correlation with endoscopic inflammation score (**Fig 6D**). This indicates that choline may serve as a fecal marker of inflammation.

The observed trends in metabolite levels do not reflect those observed in tissues, most likely because the fecal metabolome results from a complex interplay between gut microflora metabolism and host metabolism. Inflammation causes an imbalance in the gut microbiota leading to changes that are still difficult to interpret in terms of biochemical pathways, but are extremely valuable at the diagnostic level as they contain a strong and distinctive metabolomic signature already at the pre-symptomatic state.

## Conclusions

Inflammation is known to be associated with hypoxia due to the extensive activity of the immune system. Our analysis provides, for the first time, a system-level view of how this hypoxia affects the respiratory state of the whole tissue – shifting it to anaerobic respiration, as evident by the combined metabolomic and proteomic analyses (**Fig 4**). Most remarkably, however, the metabolic profile of the pre-symptomatic states is distinct compared to ill, and most importantly, to healthy WT samples. A characteristic metabolomic signature of inflammation and pre-clinical states is evident also at the fecal levels.

The two pre-symptomatic states are also very similar to each other, as revealed by testing day 4 samples on the PCA-CA model built with the other four groups. Therefore, it is apparent that though at the tissue level no macroscopic or histological signs of inflammation can be detected, the metabolism of the tissue already begins to shift, as part of the early molecular events involved in the pathology. In addition, the similarity between the two types of pre-symptomatic states, preceding either chronic or acute inflammation, indicates that metabolically, both types of inflammation pass through the same metabolic-state.

The diagnostic potential of our untargeted metabolic approach is most pronounced in the fecal sample analysis, giving rise to very high discrimination accuracy already in pre-symptomatic states with low invasiveness. Specifically, choline seems to have a very high correlation with inflammation score, which is in line with a previous study showing a similar pattern in pediatric IBD patients (Kolho et al, 2016). The measured fecal metabolome does not reflect the trends observed in tissues, reflecting the contribution of the commensal microbiota in these samples, and therefore the profound differences among states may stem from the differences in microbial diversity that are known to be associated with IBD (Bibiloni & Schiffrin, 2010; Mottawea et al, 2016; Ott & Schreiber, 2006). This is also supported by a report on changes in fatty acid composition, coming from the microbiota, in fecal samples from IBD patients (De Preter et al, 2015). Therefore, global fecal metabolomics has potential for early diagnosis, as pre-symptomatic states have their own distinct profile.

In conclusion, metabolism is profoundly affected by intestinal inflammation, from an early stage in the pathogenic process. Pre-clinical metabolic shifts can be detected on different levels of analysis – host tissue and fecal. These early metabolic signatures have a predictive potential that will aid disease management and prediction of flares.

## Materials and Methods

### Key Resources Table

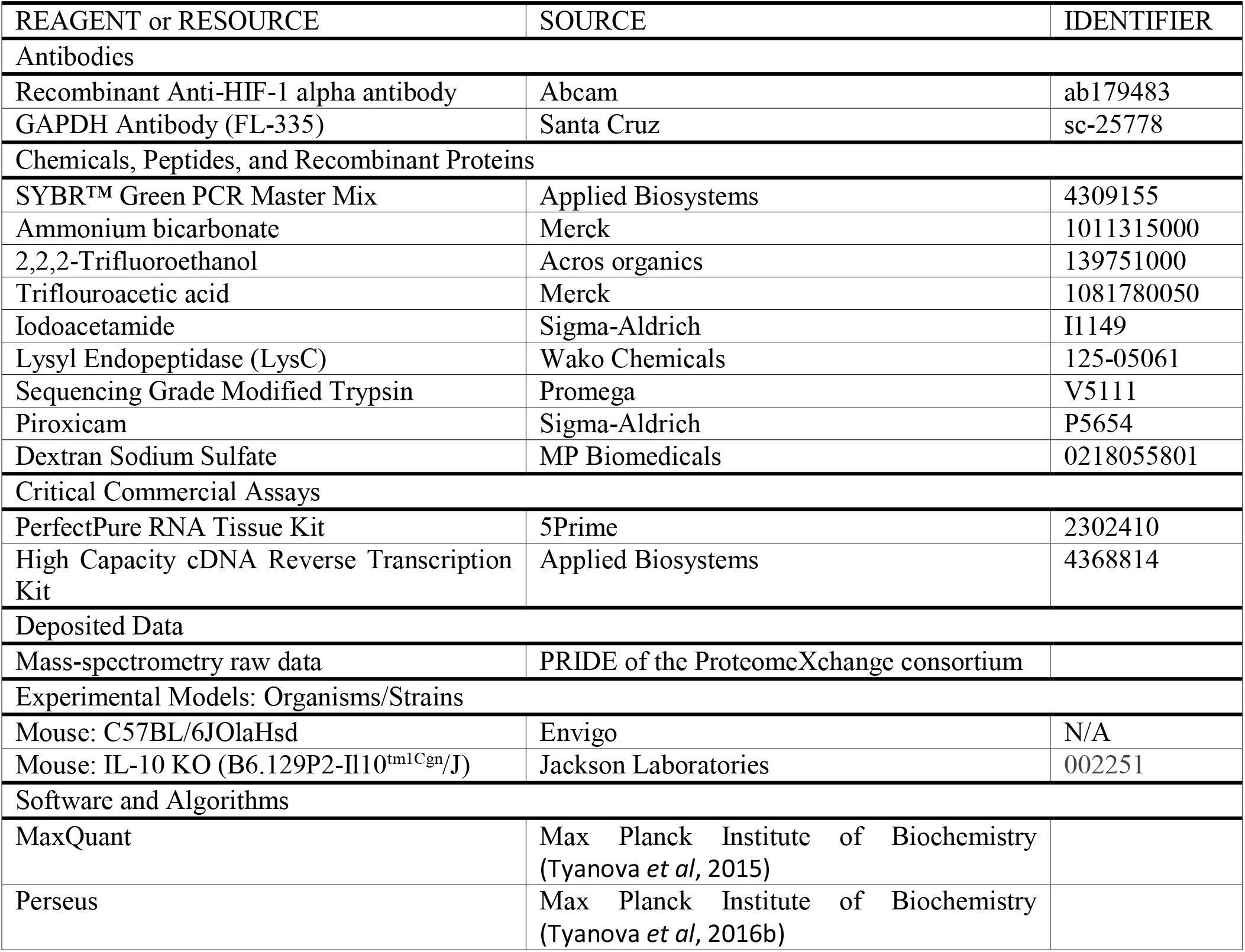

### Contact for Reagent and Resource Sharing

Further information and requests for reagents may be directed to, and will be fulfilled by the corresponding author Irit Sagi.

### Animals

Seven-week old C57BL/6 male mice were purchased from Envigo and were allowed to adapt for one week before experimental procedure. IL-10^−/−^ C57BL/6 mice from Jackson Laboratories were inbred at the Weizmann Institute of Science, and experiments were performed on six to eight-week old male mice. All experiments and procedures were approved by the Weizmann Institute of Science animal care and use committees, protocol numbers 02230413-2, 06481012-1.

### Intestinal inflammation induction and evaluation

Acute colonic inflammation was induced by administration of 1.25% DSS (MP Biomedicals LLC) in drinking water of C57BL/6 mice for seven days (Okayasu *et al*, 1990). Chronic colonic inflammation was accelerated and synchronized by peroral administration of 200ppm piroxicam (Sigma-Aldrich ltd.) to IL-10^−/−^ C57BL/6 mice via supplementation in normal murine chow(Berg *et al*, 2002a). Mice were weighed 3 times a week over the course of the experiment. Colitis progression was evaluated over the course of the experiment using the Karl Stortz Coloview mini endoscope system and colonic inflammation was scored (0-15) as previously described (Becker *et al*, 2006). Inflammation scores were categorized according to the following: healthy (0-4); mildly inflamed (5-7); inflamed (8-11); severely inflamed (12-15). Another form of inflammation evaluation was histological analysis by H&E staining of formalin-fixed paraffin-embedded tissues sections.

### NMR-based Metabolomics

Intact tissues were analyzed through High Resolution (HR) Magic Angle Spinning (MAS) NMR analysis (Vignoli *et al*, 2019; Beckonert *et al*, 2010). To this end, frozen colon samples were trimmed (15–20 mg) to fit HR-MAS ZrO_2_ rotor insert capacity (50 μL). Each insert was filled with ^2^H_2_O containing 5.8 mM sodium trimethylsilyl [2,2,3,3-^2^H_4_]propionate (TMSP). Inserts were covered with plug and plug-restraining screw and inserted into the 4 mm rotor for HR-MAS probe.

HR-MAS spectra were recorded with a Bruker 600 MHz spectrometer equipped with HR-MAS TXI ^1^H/^13^C/^15^N probe and magnetic field gradient along the magic-angle axis. Samples were spun at 4 MHz at 277 K. ^1^H NMR spectra were acquired with the Carr, Purcell, Meiboom, and Gill (CPMG) (Carr & Purcell, 1954) sequence using a 1D spin–echo sequence with water presaturation (cpmgpr, Bruker). 64 scans over a spectral region of 12 kHz were collected into 32 K points, giving an acquisition time of 1.36 s.

Free induction decays were multiplied by an exponential function equivalent to 1 Hz line broadening before applying Fourier transform. Transformed spectra were automatically corrected for phase and baseline distortions. Chemical shift was calibrated to the proton of signal at δ 1.48 ppm.

Fecal extracts were analyzed through in solution NMR experiments (Vignoli *et al*, 2019). Feces were mashed with 700 μL of ice-cold PBS and sonicated for 30 min to inactivate gut bacteria and achieve biochemical stability in the sample. The samples were then centrifuged at 10,000 g for 2 min, and 550–600 μL of the supernatant was transferred into a new Eppendorf tube and stored at −80°C before the analysis.

At the moment of the analysis, samples of frozen extracts were thawed at room temperature and shaken before use. A 630 μL aliquot of each sample was added to 70 μL of potassium phosphate buffer (1.5 M K_2_HPO_4_, 100% (v/v) ^2^H_2_O, 10 mM TMSP, at pH 7.4). 600 μL of each mixture were transferred into 4.25 mm NMR tubes (Bruker BioSpin srl) for analysis.

^1^H NMR spectra were acquired using a Bruker 600 MHz metabolic profiler (Bruker BioSpin) operating at 600.13 MHz proton Larmor frequency and equipped with a 5 mm TXI ^1^H-^13^C-^15^N and ^2^H-decoupling probe including a z axis gradient coil, an automatic tuning-matching (ATM) and an automatic sample changer (SampleJet). A BTO 2000 thermocouple served for temperature stabilization at the level of approximately 0.1 K at the sample. Before measurement, samples were kept for at least 5 minutes inside the NMR probehead for temperature equilibration (300 K). For each sample, a ^1^H NMR spectrum was acquired with the CPMG (Carr & Purcell, 1954) sequence using a 1D spin–echo sequence with water presaturation. 256 scans over a spectral region of 12 kHz were collected into 73 K points, giving an acquisition time of 3.06 s.

The raw data were multiplied by a 0.3 Hz exponential line broadening before Fourier transformation into 128 K points. Chemical shift was referenced to the signal of TMSP at δ 0.00 ppm.

All the spectra were binned for the subsequent multivariate statistical analysis. Binning is a means to reduce the number of total variables and to compensate for small shift in the signals, making the analyses more robust and reproducible.

In particular, each spectrum of tissues was segmented in the region between 4.85–0.2 ppm into 0.01 ppm chemical shift bins, and the corresponding spectral areas were integrated using the AMIX software (Bruker). The area of each bin was normalized to the total spectral area, calculated with exclusion of the lipid regions 2.88–2.72, 2.32–2.22, 2.12–1.98, 1.70–1.50, 1.46–1.22, and 0.96–0.76 ppm.

Each spectrum of fecal extract was segmented in the region between 10.0 - 0.2 ppm into 0.02 ppm chemical shift bins. The spectral region between 6.0 - 4.2 ppm, containing the water signal, was discarded. Finally, total area normation was carried out.

The metabolites, whose peaks in the NMR spectra were well defined and resolved, were assigned. Signal identification was achieved using a library of NMR spectra of pure organic compounds, public databases (such as HMBD, Human Metabolic Database) (Wishart *et al*, 2012) storing reference NMR spectra of metabolites, spiking NMR experiments and literature data. The relative concentrations of the various metabolites in the different spectra were calculated by integrating the signal area(Vignoli *et al*, 2019; Wishart, 2008).

### Proteomics by liquid chromatography–tandem mass spectrometry (LC-MS/MS) analysis

Tissue slices from colons of WT or IL-10^−/−^ mice at different stages of the inflammation models were immersed in a solution containing 50% trifluoroethanol (TFE), 25mM ammonium bicarbonate (Sigma Aldrich) and 15mM dithiothreitol (DTT) and coarsely homogenized by repeated cycles of boiling, freezing and sonication. Subsequently, samples were shaken at 30°C, 1400 rpm for 30 min, followed by the addition of iodoacetamide (Sigma Aldrich) to a final concentration of 25 mM and further shaking for 30 min at 30°C and 1400 rpm. TFE was then diluted to 25% with 50 mM ABC, LysC (Wako Chemicals, 1:100 lysC:protein ratio) and sequencing-grade modified trypsin (Promega, 1:50 trypsin:protein ratio) was added and incubated overnight at room temperature. On the following morning, more trypsin was added (1:80 trypsin: protein ratio) for 4 hours. Peptide mixtures were purified on C18 stage tips. Eluted peptides were loaded onto a 50 cm long EASY-spray reverse phase column and analyzed on an EASY- nLC- 1000 HPLC system (Thermo Scientific) coupled to a Q-Exactive Plus MS (Thermo Scientific). Peptides were separated over 240 minutes with a gradient of 5−28 % buffer B (80% acetonitrile and 0.1% formic acid). One full MS scan was acquired at a resolution of 70,000 and each full scan was followed by the selection of 10 most intense ions (Data dependent Top 10 method) at a resolution of 17,500 for MS2 fragmentation.

Raw MS data was analyzed with MaxQuant software (Cox & Mann, 2008; Tyanova *et al*, 2016a) (version 1.5.2.18) with the built-in Andromeda search engine(Cox *et al*, 2011) by searching against the mouse reference proteome (UniprotKB,Nov2014). Enzyme specificity was set to trypsin cleavage after lysine and arginine and up to two miscleavages were allowed. The minimum peptide length was set to seven amino acids. Acetylation of protein N termini, deamidation of asparagine and glutamine, and oxidation of methionine were set as variable modifications. Carbamidomethylation of cysteine was set as a fixed modification. Protein identifications were sorted using a target-decoy approach at a false discovery rate (FDR) of 1% at the peptide and protein levels. Relative, label-free quantification of proteins was performed using the MaxLFQ algorithm integrated into MaxQuant environment with minimum ratio count of two(Cox *et al*, 2014).

Bioinformatics analysis was performed with the Perseus program version 1.5.1.4 (Tyanova *et al*, 2016b). The proteomic data was first filtered to remove the potential contaminants, proteins only identified by their modification site and reverse proteins. Next, the intensity values were log2 transformed and data was filtered to have at least three valid values in each group. Missing values were imputed based on normal distribution. Proteins appearing in the Human Metabolic Atlas (http://www.metabolicatlas.org/) were chosen for analysis.

### Western blot analysis

Tissues were manually homogenized in RIPA buffer (20 mM Tris pH 7.4, 137 mM, NaCl, 10% glycerol, 0.1% SDS, 0.5% deoxycholate, 1% Triton, 2 mM EDTA, 1 mM phenylmethylsulfonyl fluoride, 20 μM leupeptin, 1 tablet/100 ml of cOmplete™ Protease Inhibitor Cocktail (Roche)). Western blot analysis was performed using anti-HIF1α antibody (Abcam), for detection of the protein at 110-120 kD. Anti-GAPDH (Santa Cruz) was used as the loading control and the protein was detected at 35 kD. 3-4 samples from different mice were run for each tissue state, and quantification was carried out using ImageJ.

### Statistical analysis

Multivariate statistical analyses were performed both on binned NMR-spectra, for metabolomics, and on metabolism-associated proteins, for proteomics.

Various kinds of multivariate statistical techniques were applied using R 3.0.2 in house scripts. Principal Component Analysis (PCA) was used to obtain a preliminary outlook of the data (visualization in a reduced space, clusters detection, screening for outliers). Canonical analysis (CA) was used in combination with PCA to increase the supervised separation of the analyzed groups. The global accuracy for classification was assessed by means of a Monte Carlo cross-validation scheme. Accordingly, each dataset was randomly divided into a training set (90% of the data) and a test set (10% of the data). The training set was used to build the model, whereas the test set was used to validate its discriminant and predictive power; this operation was repeated 500 times. Accuracy, specificity and sensitivity were estimated according to standard definitions.

Student’s t-test with false discovery rate (FDR) correction (Tusher et al., 2001), were used for the determination of the meaningful metabolites and differentially abundant proteins.

### Data and Software Availability

The MS proteomics data have been deposited to the ProteomeXchange Consortium via the PRIDE partner repository with the dataset identifier PXD004740 and can be viewed by logging in with the following details: Username; Password: hKx9zFSu.

The NMR data have been deposited to the MetaboLights database (www.ebi.ac.uk/metabolights) with the accession number MTBLS1496.

## Author Contributions

Conceptualization, ES and IS; Methodology, ES and VG; Formal Analysis, ES and VG; Resources, PT, CL and IS; Writing, ES, VG, PT and IS; Visualization, ES and VG; Supervision, PT, CL and IS; Funding Acquisition, VG, PT, CL and IS.

## Acknowledgements

PT, VG and CL acknowledge the support and the use of resources of Instruct-ERIC, a Landmark ESFRI project, and specifically the CERM/CIRMMP Italy Centre. VG thanks EMBO for a Short Term Fellowship (ASTF: 135 - 2016).

