## Supplementary Information for "Integrated metabolomics and proteomics of symptomatic and early pre-symptomatic states of colitis"

#Authors contributed equally

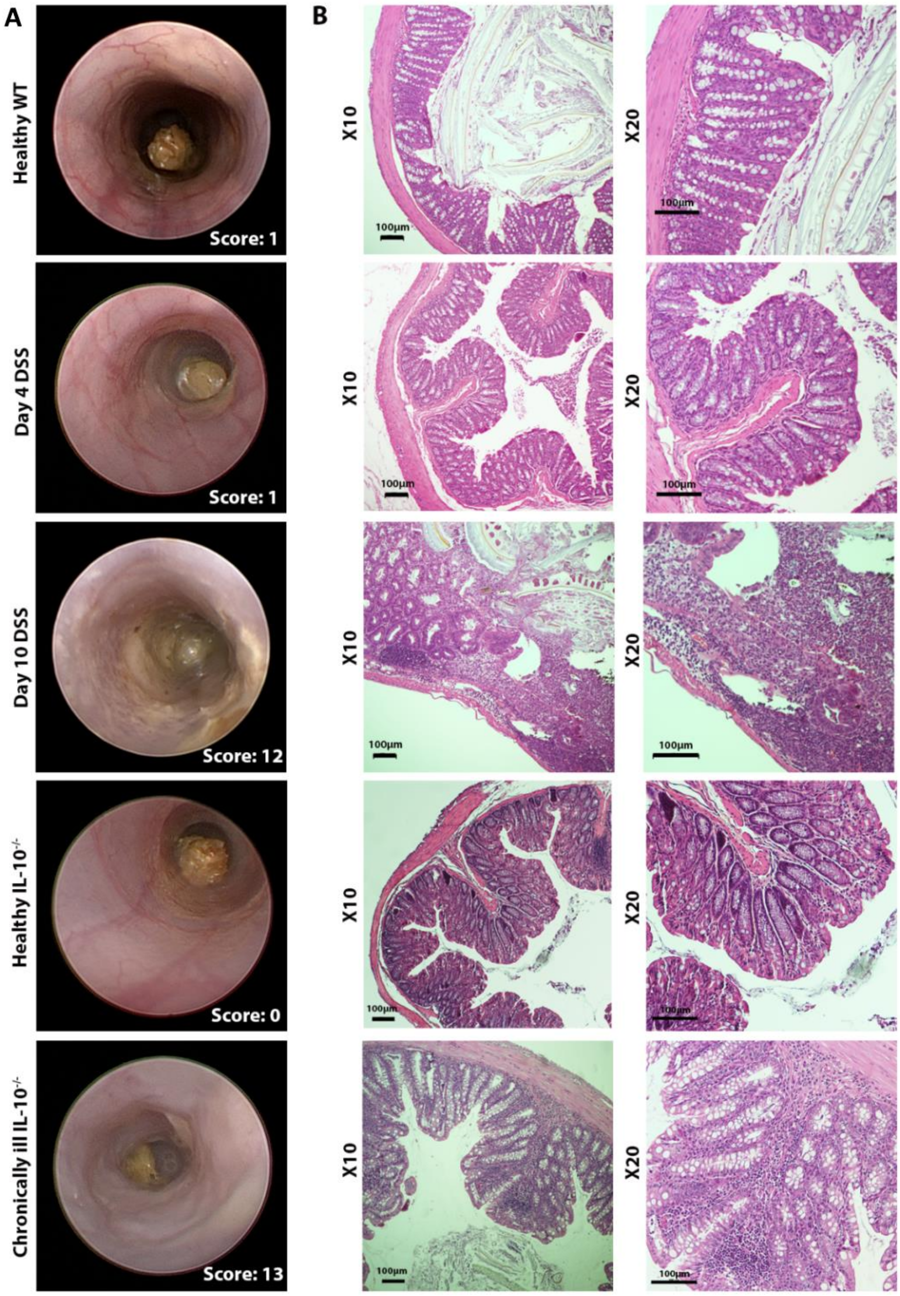


***Figure S1***. *A. Endoscopic images of colon in live mice at the indicated states. Each mouse is given a score based on five parameters (colon transparency, changes in vascular pattern, feces consistency, fibrin deposition and granularity of mucosal surface) was given: 0-4: healthy; 5-7: mildly inflamed; 8-11: inflamed; 12-15: severely inflamed. B. Hematoxylin and eosin-stained colonic sections of mice at the indicated states. Immune cell infiltration and mucosal damage is not evident on day 4 of the acute model, but is substantial on day 10. A large amount of immune cell infiltrate into the mucosa evident in the chronic IL-10-/- model.*

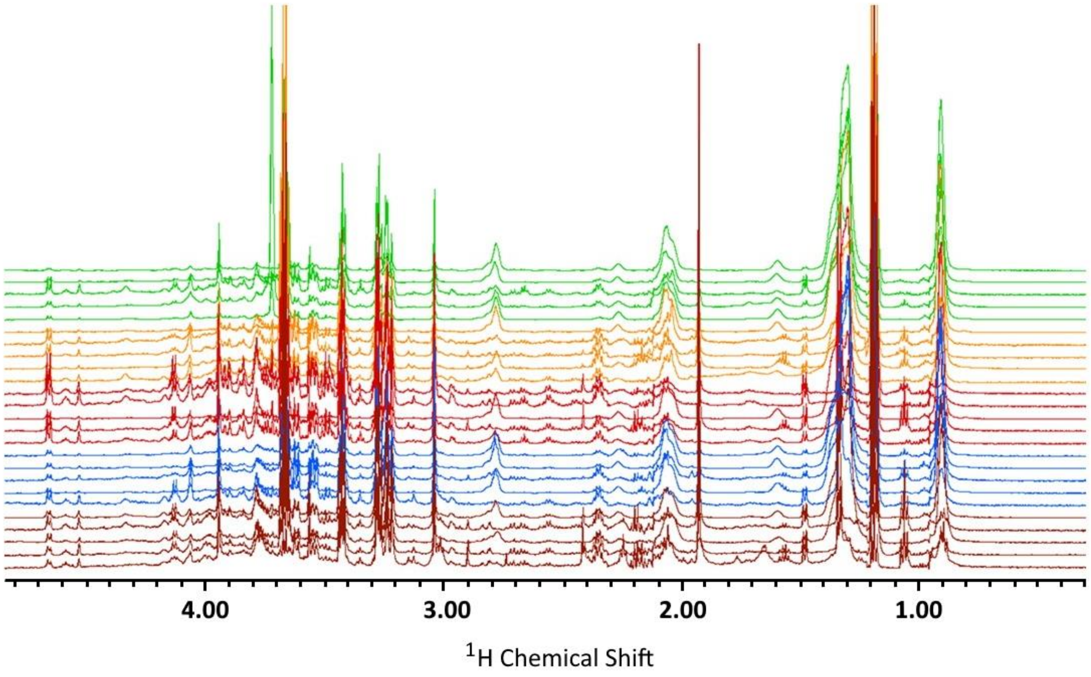


***Figure S2***. *HR-MAS* $^{1}$*H NMR spectra of colon tissues. The spectra of five samples per group are reported. Green traces, healthy WT (WT-HC) mice; orange traces, mice on day 4 of the DSS model (WT-D4); red traces, acutely ill mice on day 10 of the DSS model (WT-D10);blue dots, healthy IL-10-/- mice (IL10-HC); brown traces, chronically ill IL-10-/- mice (IL10-ILL).*

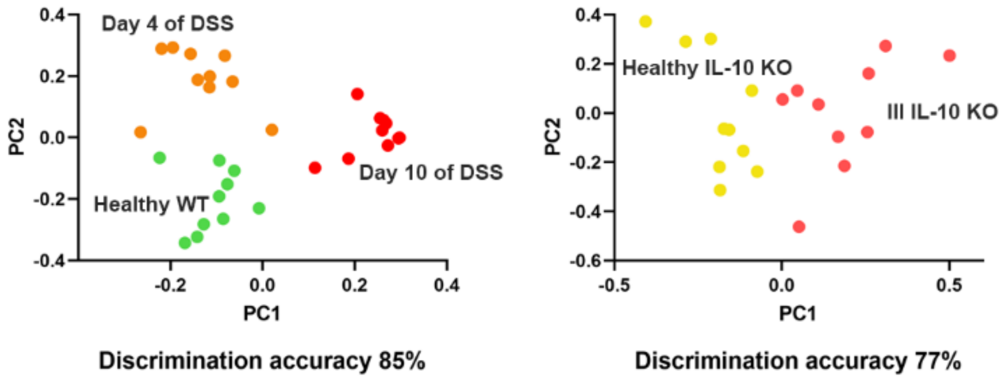


***Figure S3.*** *Untargeted NMR-based metabolomic profiles of the two murine models: DSS- model (left) and IL10$^{-/-}$ model (right). The PCA-CA score plots with the respective discrimination accuracy values are reported. In both the PCA-CA score plots, each dot represents a different sample and each color a different group of mice: green dots, healthy WT (WT-HC) mice; orange dots, mice on day 4 of the DSS model (WT-D4); yellow dots, healthy IL-10-/- mice (IL10-HC); red dots, acutely ill mice on day 10 of the DSS model (WT-D10); pink dots, chronically ill IL-10-/- mice (IL10-ILL).*

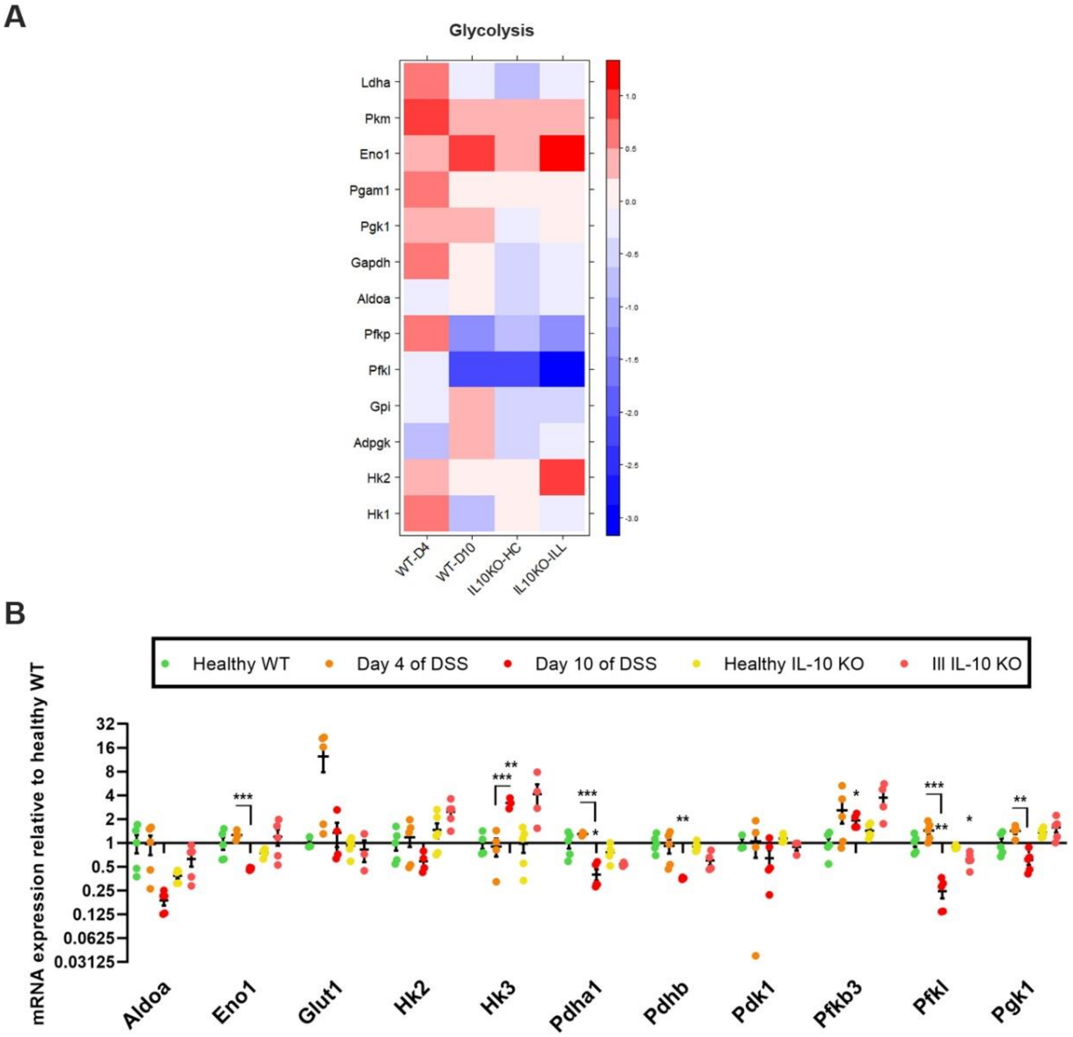


***Figure S4.*** *(A) Heat maps for relative mean protein abundance (in log2(LFQ) compared to healthy WT) of proteins involved in glycolysis. (B) Box plots of relative levels of mRNA data of glycolytic proteins obtained by real-time PCR across the five colitis stages. Green dots, healthy WT (WT-HC) mice; orange dots, mice on day 4 of the DSS model (WT-D4); yellow dots, healthy IL-10-/- mice (IL10-HC); red dots, acutely ill mice on day 10 of the DSS model (WT-D10); pink dots, chronically ill IL-10-/- mice (IL10-ILL).*

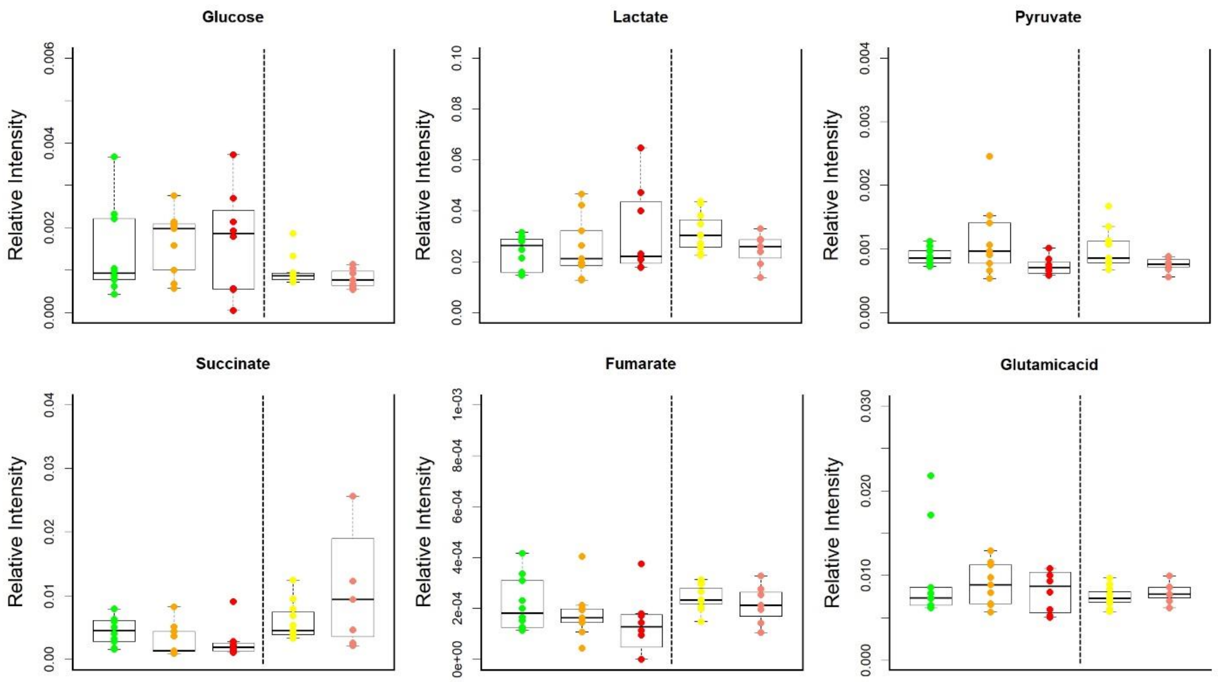


***Figure S5.*** *Box plots of relative levels of metabolites involved in glucose metabolism in fecal extracts. Green dots, healthy WT (WT-HC) mice; orange dots, mice on day 4 of the DSS model (WT-D4); yellow dots, healthy IL-10-/- mice (IL10-HC); red dots, acutely ill mice on day 10 of the DSS model (WT-D10); pink dots, chronically ill IL-10-/- mice (IL10-ILL).*

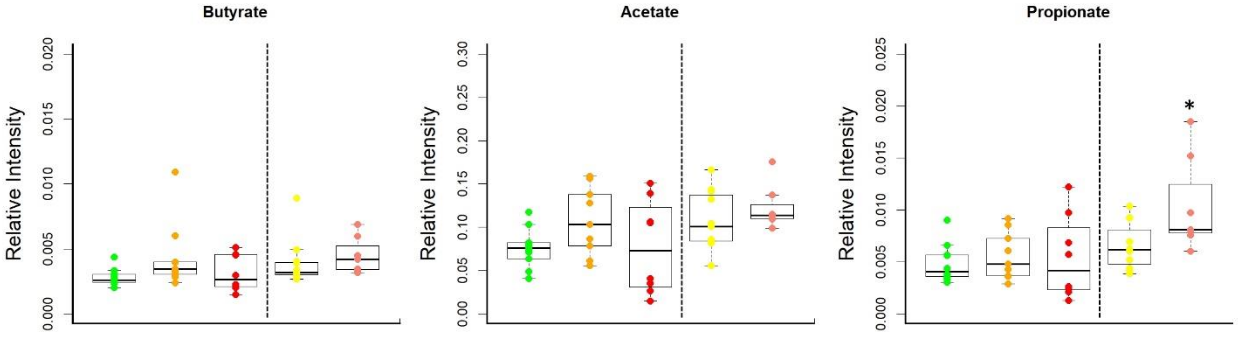


***Figure S6.*** *Box plots of relative levels of butyrate, acetate and propionate in fecal extracts. Green dots, healthy WT (WT-HC) mice; orange dots, mice on day 4 of the DSS model (WT-D4); yellow dots, healthy IL-10-/- mice (IL10-HC); red dots, acutely ill mice on day 10 of the DSS model (WT-D10); pink dots, chronically ill IL-10-/- mice (IL10-ILL). Asterisks indicate significance relative to corresponding healthy state (either WT or IL10KO), unless otherwise indicated by bars. *p<0.05.*

***Table S1***: *List of the metabolites assigned and analyzed in colon tissues. The Human Metabolome Database (HMDB) compound ID of each metabolite is reported.*

| **Metabolite** | **HMDB ID** |
|---|---|
| Acetate | HMDB00042 |
| L-Alanine | HMDB00161 |
| Ascorbic acid | HMDB00044 |
| L-Aspartic acid | HMDB00191 |
| Choline | HMDB00097 |
| Creatine | HMDB00064 |
| Ethanol | HMDB00108 |
| Fumarate | HMDB00134 |
| D-Glucose | HMDB00122 |
| L-Glutamic acid | HMDB00148 |
| Glutathione | HMDB00125 |
| Glycine | HMDB00123 |
| Hypotaurine | HMDB00965 |
| L-Isoleucine | HMDB00172 |
| Lactate | HMDB00190 |
| Malonate | HMDB00691 |
| Myo-inositol | HMDB00211 |
| Propionate | HMDB00237 |
| Succinate | HMDB00254 |
| Taurine | HMDB00251 |
| Trimethylamine | HMDB00906 |
| L-Valine | HMDB00883 |

***Table S2****: Comparison of protein levels involved in different metabolic processes among the different colitis states. The P-values are corrected for FDR formula.*

| Glycolysis | | | | | | |
|---|---|---|---|---|---|---|
| Protein | WT-HC vs WT-D10 | WT-HC vs WT-D4 | WT-D4 vs WT-D10 | IL10$^{-/-}$HC vs IL10$^{-/-}$ILL | WT-HC vs IL10$^{-/}$-HC | WT-D10 Vs IL10$^{-/-}$ILL |
| Hk1 | 0.072 | 0.7621 | 0.0153 | 0.6062 | 0.9022 | 0.358 |
| Hk2 | 0.5995 | 0.7621 | 0.1594 | 0.1011 | 0.9022 | 0.2844 |
| Adpgk | 0.2395 | 0.8288 | 0.0446 | 0.7032 | 0.7944 | 0.6502 |
| Gpi | 0.542 | 0.9021 | 0.3163 | 0.7454 | 0.5241 | 0.2844 |
| Pfkl | 0.0116 | 0.8728 | 0.0191 | 0.7454 | 0.4156 | 0.0162 |
| Pfkp | 0.2871 | 0.2444 | 0.0039 | 0.7454 | 0.6125 | 0.2844 |
| Aldoa | 0.9473 | 0.8728 | 0.903 | 0.7454 | 0.4156 | 0.2844 |
| Gapdh | 0.5995 | 0.2444 | 0.6451 | 0.7454 | 0.5241 | 0.4264 |
| Pgk1 | 0.5995 | 0.7621 | 0.9387 | 0.7032 | 0.6125 | 0.9034 |
| Pgam1 | 0.5995 | 0.1703 | 0.1663 | 0.7454 | 0.6125 | 0.6502 |
| Eno1 | 0.2395 | 0.7621 | 0.4747 | 0.1011 | 0.7944 | 0.1699 |
| Pkm | 0.5995 | 0.1017 | 0.1663 | 0.7454 | 0.6125 | 0.2844 |
| Ldha | 0.8732 | 0.8288 | 0.5802 | 0.7032 | 0.4156 | 0.5825 |

| TCA Cycle | | | | | | |
|---|---|---|---|---|---|---|
| Protein | WT-HC vs WT-D10 | WT-HC vs WT-D4 | WT-D4 vs WT-D10 | IL10$^{-/-}$HC vs IL10$^{-/-}$ILL | WT-HC vs IL10$^{-/-}$HC | WT-D10 Vs IL10$^{-/-}$ILL |
| Pdha1 | 0.079 | 0.814 | 0.093 | 0.225 | 0.139 | 0.010 |
| Pdhb | 0.893 | 0.814 | 0.853 | 0.267 | 0.887 | 0.228 |
| Dlat | 0.026 | 0.702 | 0.163 | 0.264 | 0.140 | 0.002 |
| Dld | 0.030 | 0.702 | 0.091 | 0.108 | 0.140 | 0.006 |
| Pc | 0.005 | 0.702 | 0.00001 | 0.0003 | 0.825 | 0.013 |
| Acly | 0.0002 | 0.164 | 0.00001 | 0.0003 | 0.083 | 0.173 |
| Aco1 | 0.344 | 0.816 | 0.431 | 0.611 | 0.825 | 0.522 |
| Aco2 | 0.034 | 0.702 | 0.091 | 0.006 | 0.207 | 0.002 |
| Idh1 | 0.214 | 0.898 | 0.339 | 0.718 | 0.140 | 0.062 |
| Idh2 | 0.309 | 0.702 | 0.946 | 0.067 | 0.407 | 0.002 |
| Idh3g | 0.181 | 0.702 | 0.557 | 0.085 | 0.405 | 0.225 |
| Idh3a | 0.157 | 0.702 | 0.340 | 0.009 | 0.248 | 0.017 |
| Idh3b | 0.038 | 0.702 | 0.102 | 0.31 | 0.108 | 0.008 |
| Ogdh | 0.030 | 0.702 | 0.002 | 0.002 | 0.297 | 0.002 |
| Suclg1 | 0.030 | 0.449 | 0.114 | 0.023 | 0.990 | 0.030 |
| Suclg2 | 0.038 | 0.816 | 0.082 | 0.000 | 0.207 | 0.002 |
| Sdha | 0.004 | 0.814 | 0.002 | 0.026 | 0.248 | 0.004 |
| Sdhb | 0.030 | 0.702 | 0.076 | 0.108 | 0.134 | 0.004 |
| Fh | 0.118 | 0.778 | 0.393 | 0.283 | 0.108 | 0.002 |
| Mdh1 | 0.568 | 0.816 | 0.435 | 0.173 | 0.299 | 0.055 |
| Mdh2 | 0.568 | 0.164 | 0.114 | 0.225 | 0.140 | 0.014 |
| Pck2 | 0.157 | 0.245 | 0.435 | 0.108 | 0.134 | 0.381 |

| β-Oxidation | | | | | | |
|---|---|---|---|---|---|---|
| Protein | WT-HC vs WT-D10 | WT-HC vs WT-D4 | WT-D4 vs WT-D10 | $IL10^{-/-}$HC vs $IL10^{-/-}$ILL | WT-HC vs $IL10^{-/-}$-HC | WT-D10 Vs $IL10^{-/-}$ILL |
| Acsl1 | 0.135 | 0.581 | 0.152 | 0.721 | 0.53 | 0.284 |
| Acsl5 | 0.956 | 0.581 | 0.479 | 0.761 | 0.492 | 0.545 |
| Cpt1a | 0.247 | 0.581 | 0.013 | 0.017 | 0.79 | 0.074 |
| Cpt2 | 0.144 | 0.59 | 0.27 | 0.088 | 0.15 | 0.003 |
| Acadm | 0.045 | 0.581 | 0.094 | 0.009 | 0.404 | 0.002 |
| Acadvl | 0.309 | 0.581 | 0.7 | 0.22 | 0.174 | 0.002 |
| Acadl | 0.144 | 0.581 | 0.087 | 0.22 | 0.592 | 0.434 |
| Acads | 0.048 | 0.517 | 0.088 | 0.818 | 0.001 | 0.00009 |
| Gcdh | 0.045 | 0.581 | 0.061 | 0.017 | 0.457 | 0.03 |
| Ehhadh | 0.045 | 0.581 | 0.05 | 0.237 | 0.662 | 0.329 |
| Echs1 | 0.073 | 0.609 | 0.05 | 0.274 | 0.241 | 0.036 |
| Hadha | 0.047 | 0.581 | 0.002 | 0.004 | 0.731 | 0.027 |
| Hadh | 0.158 | 0.581 | 0.034 | 0.034 | 0.621 | 0.017 |
| Acaa1 | 0.554 | 0.581 | 0.13 | 0.314 | 0.79 | 0.545 |
| Acaa2 | 0.032 | 0.59 | 0.01 | 0.004 | 0.031 | 0.00008 |
| Hadhb | 0.045 | 0.971 | 0.034 | 0.75 | 0.196 | 0.009 |
| Acat1 | 0.032 | 0.581 | 0.06 | 0.009 | 0.119 | 0.0004 |
| Acat2 | 0.994 | 0.581 | 0.27 | 0.036 | 0.483 | 0.027 |
| Eci1 | 0.057 | 0.517 | 0.497 | 0.75 | 0.031 | 0.002 |
| Eci2 | 0.095 | 0.971 | 0.034 | 0.342 | 0.592 | 0.048 |
| Adh1 | 0.002 | 0.25 | 0.00001 | 0.004 | 0.548 | 0.017 |
| Aldh2 | 0.095 | 0.581 | 0.006 | 0.004 | 0.731 | 0.004 |
| Aldh1b1 | 0.045 | 0.581 | 0.06 | 0.004 | 0.075 | 0.00008 |
| Aldh3a2 | 0.095 | 0.682 | 0.272 | 0.761 | 0.353 | 0.071 |

| BCAA Degradation | | | | | | |
|---|---|---|---|---|---|---|
| Protein | WT-HC vs WT-D10 | WT-HC vs WT-D4 | WT-D4 vs WT-D10 | $IL10^{-/-}$HC vs $IL10^{-/-}$ILL | WT-HC vs $IL10^{-/-}$-HC | WT-D10 Vs $IL10^{-/-}$ILL |
| Bcat2 | 0.078 | 0.009 | 0.462 | 0.241 | 0.001 | 0.009 |
| Bckdha | 0.029 | 0.609 | 0.138 | 0.013 | 0.938 | 0.028 |
| Dld | 0.033 | 0.609 | 0.065 | 0.138 | 0.165 | 0.008 |
| Dbt | 0.038 | 0.609 | 0.136 | 0.095 | 0.102 | 0.009 |
| Acadm | 0.035 | 0.609 | 0.102 | 0.01 | 0.328 | 0.001 |
| Acads | 0.038 | 0.35 | 0.095 | 0.886 | 0.001 | 0.000005 |
| Ehhadh | 0.033 | 0.609 | 0.055 | 0.241 | 0.683 | 0.312 |
| Echs1 | 0.059 | 0.692 | 0.055 | 0.279 | 0.214 | 0.035 |
| Hadha | 0.038 | 0.609 | 0.002 | 0.005 | 0.757 | 0.027 |
| Mccc2 | 0.052 | 0.609 | 0.122 | 0.863 | 0.328 | 0.5 |
| Hibch | 0.321 | 0.991 | 0.469 | 0.46 | 0.343 | 0.035 |
| Hadh | 0.148 | 0.609 | 0.042 | 0.041 | 0.64 | 0.017 |
| Hmgcl | 0.414 | 0.35 | 0.567 | 0.871 | 0.09 | 0.01 |
| Oxct1 | 0.078 | 0.802 | 0.055 | 0.188 | 0.938 | 0.096 |
| Acat1 | 0.026 | 0.609 | 0.065 | 0.01 | 0.09 | 0.0004 |
| Acat2 | 0.994 | 0.609 | 0.266 | 0.043 | 0.378 | 0.027 |
| Hibadh | 0.099 | 0.991 | 0.118 | 0.915 | 0.378 | 0.284 |

| | | | | | | |
|---|---|---|---|---|---|---|
| Acaa1 | 0.528 | 0.609 | 0.126 | 0.322 | 0.855 | 0.545 |
| Acaa2 | 0.026 | 0.653 | 0.008 | 0.005 | 0.023 | 0.00006 |
| Aldh6a1 | 0.027 | 0.695 | 0.042 | 0.241 | 0.425 | 0.049 |
| Aldh2 | 0.078 | 0.609 | 0.005 | 0.005 | 0.757 | 0.006 |
| Aldh3a2 | 0.078 | 0.739 | 0.269 | 0.863 | 0.294 | 0.069 |
| Aldh1b1 | 0.033 | 0.609 | 0.065 | 0.005 | 0.065 | 0.00006 |
| Pcca | 0.0003 | 0.008 | 0.002 | 0.005 | 0.008 | 0.000004 |
| Pccb | 0.026 | 0.886 | 0.002 | 0.915 | 0.078 | 0.017 |
| Mut | 0.033 | 0.609 | 0.073 | 0.624 | 0.282 | 0.121 |
| **Oxidative Phosphorylation** | | | | | | |
| Protein | WT-HC vs WT-D10 | WT-HC vs WT-D4 | WT-D4 vs WT-D10 | IL10$^{-/-}$HC vs IL10$^{-/-}$ILL | WT-HC vs IL10$^{-/}$-HC | WT-D10 Vs IL10$^{-/-}$ILL |
| Ndufs1 | 0.034 | 0.981 | 0.003 | 0.001 | 0.961 | 0.012 |
| Ndufs2 | 0.049 | 0.981 | 0.018 | 0.003 | 0.581 | 0.004 |
| Ndufs3 | 0.088 | 0.981 | 0.038 | 0.003 | 0.581 | 0.001 |
| Ndufs5 | 0.049 | 0.981 | 0.117 | 0.005 | 0.581 | 0.012 |
| Ndufs6 | 0.02 | 0.631 | 0.071 | 0.005 | 0.839 | 0.004 |
| Ndufs7 | 0.064 | 0.981 | 0.046 | 0.022 | 0.321 | 0.445 |
| Ndufs8 | 0.133 | 0.981 | 0.149 | 0.001 | 0.839 | 0.004 |
| Ndufv1 | 0.025 | 0.981 | 0.001 | 0.002 | 0.839 | 0.006 |
| Ndufv2 | 0.049 | 0.981 | 0.046 | 0.804 | 0.321 | 0.049 |
| Ndufv3 | 0.089 | 0.189 | 0.379 | 0.804 | 0.635 | 0.098 |
| Ndufa2 | 0.049 | 0.981 | 0.046 | 0.005 | 0.581 | 0.004 |
| Ndufa4 | 0.723 | 0.89 | 0.228 | 0.987 | 0.321 | 0.068 |
| Ndufa5 | 0.049 | 0.981 | 0.054 | 0.002 | 0.581 | 0.001 |
| Ndufa6 | 0.04 | 0.981 | 0.019 | 0.11 | 0.77 | 0.066 |
| Ndufa7 | 0.02 | 0.631 | 0.046 | 0.072 | 0.77 | 0.013 |
| Ndufa8 | 0.053 | 0.631 | 0.267 | 0.265 | 0.465 | 0.397 |
| Ndufa9 | 0.036 | 0.981 | 0.07 | 0.005 | 0.635 | 0.006 |
| Ndufa10 | 0.057 | 0.991 | 0.046 | 0.081 | 0.839 | 0.018 |
| Ndufa12 | 0.064 | 0.981 | 0.117 | 0.012 | 0.839 | 0.006 |
| Ndufa13 | 0.049 | 0.981 | 0.046 | 0.176 | 0.581 | 0.012 |
| Ndufb3 | 0.049 | 0.981 | 0.018 | 0.603 | 0.581 | 0.128 |
| Ndufb4 | 0.049 | 0.189 | 0.881 | 0.055 | 0.839 | 0.037 |
| Ndufb5 | 0.794 | 0.847 | 0.067 | 0.804 | 0.839 | 0.916 |
| Ndufb6 | 0.088 | 0.981 | 0.089 | 0.11 | 0.839 | 0.132 |
| Ndufb7 | 0.133 | 0.981 | 0.117 | 0.352 | 0.787 | 0.143 |
| Ndufb8 | 0.089 | 0.847 | 0.379 | 0.184 | 0.581 | 0.035 |
| Ndufb9 | 0.293 | 0.981 | 0.379 | 0.055 | 0.77 | 0.226 |
| Ndufb10 | 0.558 | 0.981 | 0.561 | 0.933 | 0.839 | 0.889 |
| Ndufb11 | 0.33 | 0.981 | 0.267 | 0.804 | 0.581 | 0.354 |
| Ndufc2 | 0.21 | 0.981 | 0.267 | 0.286 | 0.839 | 0.132 |
| Sdha | 0.013 | 0.891 | 0.014 | 0.058 | 0.324 | 0.006 |
| Sdhb | 0.101 | 0.891 | 0.16 | 0.139 | 0.165 | 0.006 |
| Cyc1 | 0.407 | 0.891 | 0.225 | 0.611 | 0.079 | 0.006 |
| Uqcrb | 0.407 | 0.96 | 0.309 | 0.022 | 0.368 | 0.008 |
| Uqcrq | 0.874 | 0.891 | 0.894 | 0.956 | 0.863 | 0.893 |

| | | | | | | |
|---|---|---|---|---|---|---|
| Uqcrfs1 | 0.407 | 0.891 | 0.307 | 0.097 | 0.393 | 0.008 |
| Uqcrc1 | 0.278 | 0.972 | 0.225 | 0.096 | 0.212 | 0.006 |
| Uqcrc2 | 0.183 | 0.462 | 0.894 | 0.197 | 0.344 | 0.009 |
| Uqcr10 | 0.768 | 0.891 | 0.941 | 0.594 | 0.579 | 0.189 |
| Uqcrh | 0.801 | 0.972 | 0.894 | 0.608 | 0.682 | 0.29 |
| Cox4i1 | 0.407 | 0.462 | 0.894 | 0.29 | 0.165 | 0.005 |
| Cox5a | 0.497 | 0.96 | 0.307 | 0.056 | 0.343 | 0.016 |
| Cox7c | 0.684 | 0.891 | 0.506 | 0.13 | 0.579 | 0.298 |
| Cox5b | 0.594 | 0.891 | 0.894 | 0.96 | 0.165 | 0.172 |
| Cox6a1 | 0.826 | 0.891 | 0.894 | 0.956 | 0.958 | 0.969 |
| Cox7a2 | 0.448 | 0.891 | 0.225 | 0.022 | 0.165 | 0.004 |
| Cox6b1 | 0.465 | 0.891 | 0.941 | 0.608 | 0.274 | 0.341 |
| Cox6c | 0.368 | 0.891 | 0.506 | 0.943 | 0.242 | 0.128 |
| Atp5a1 | 0.407 | 0.891 | 0.225 | 0.797 | 0.165 | 0.111 |
| Atp5b | 0.407 | 0.96 | 0.307 | 0.022 | 0.165 | 0.002 |
| Atp5c1 | 0.768 | 0.891 | 0.681 | 0.951 | 0.703 | 0.727 |
| Atp5f1 | 0.826 | 0.891 | 0.384 | 0.197 | 0.376 | 0.007 |
| Atp5j | 0.611 | 0.891 | 0.204 | 0.235 | 0.212 | 0.011 |
| Atp5k | 0.407 | 0.891 | 0.894 | 0.211 | 0.393 | 0.037 |
| Atp5l | 0.611 | 0.891 | 0.894 | 0.11 | 0.958 | 0.172 |
| Atp5d | 0.768 | 0.891 | 0.996 | 0.115 | 0.995 | 0.238 |
| Atp5j2 | 0.169 | 0.991 | 0.185 | 0.29 | 0.995 | 0.244 |
| Atp5h | 0.407 | 0.972 | 0.303 | 0.207 | 0.151 | 0.008 |
| Atp5o | 0.407 | 0.891 | 0.204 | 0.096 | 0.218 | 0.011 |
| Atp6v0a1 | 0.073 | 0.891 | 0.172 | 0.594 | 0.151 | 0.011 |
| Atp6v1a | 0.407 | 0.891 | 0.941 | 0.951 | 0.863 | 0.969 |
| Atp6v1e1 | 0.713 | 0.891 | 0.941 | 0.611 | 0.599 | 0.264 |
| Atp6v0d1 | 0.024 | 0.92 | 0.056 | 0.022 | 0.783 | 0.031 |
| Atp6v1b2 | 0.407 | 0.891 | 0.307 | 0.096 | 0.212 | 0.006 |
| Atp6v1f | 0.95 | 0.972 | 0.941 | 0.096 | 0.212 | 0.465 |
| Ppa1 | 0.294 | 0.891 | 0.16 | 0.096 | 0.579 | 0.024 |
| Ppa2 | 0.101 | 0.462 | 0.307 | 0.11 | 0.015 | 0.001 |

***Table S3**: List of the metabolites assigned and analyzed in fecal extracts. The Human Metabolome Database (HMDB) compound ID of each metabolite is reported.*

| | |
|---|---|
| 4-hydroxyphenylacetate | HMDB0060390 |
| Acetate | HMDB0000042 |
| Acetoin | HMDB0003243 |
| Acetone | HMDB0001659 |
| L-Alanine | HMDB0000161 |
| L-Aspartic acid | HMDB0000191 |
| Butyrate | HMDB0000039 |
| Choline | HMDB0000097 |
| Creatine | HMDB0000064 |
| Dimethylamine | HMDB0000087 |
| Ethanol | HMDB0000108 |
| Formate | HMDB0000142 |
| Fumarate | HMDB0000134 |
| D-Glucose | HMDB0000122 |
| L-Glutamic acid | HMDB0000148 |
| Glycerol | HMDB0000131 |
| Glycine | HMDB0000123 |
| L-Isoleucine | HMDB0000172 |
| Isopropanol | HMDB0000863 |
| Isovalerate | HMDB0000718 |
| Lactate | HMDB0000190 |
| Methanol | HMDB0001875 |
| L-Methionine | HMDB0000696 |
| Methylamine | HMDB0000164 |
| L-Phenylalanine | HMDB0000159 |
| Propionate | HMDB0000237 |
| Putrescine | HMDB0001414 |
| Pyruvate | HMDB0000243 |
| Succinate | HMDB0000254 |
| Taurine | HMDB0000251 |
| Trimethylamine | HMDB0000906 |
| L-Threonine | HMDB0000167 |
| L-Tyrosine | HMDB0000158 |
| Uracil | HMDB0000300 |
| L-Valine | HMDB0000883 |
